# Distinct regulation of intrinsic persistent firing by cholinergic, noradrenergic and dopaminergic neuromodulations in the mouse auditory cortex

**DOI:** 10.64898/2026.05.03.722484

**Authors:** A. Tobias Price, Antonio Reboreda, Janelle Pakan, Motoharu Yoshida

## Abstract

Neuromodulators shape brain function by regulating neural activity in area- and state-dependent manners. While persistent firing (PF), a cellular correlate of short-term information retention, is strongly regulated by neuromodulation in executive and memory areas, such regulation in the sensory cortex remains less well understood. Here, using *in vitro* patch-clamp recordings, we investigated how cholinergic, noradrenergic, and dopaminergic systems regulate intrinsic PF in mouse auditory cortex (AC) neurons. We found that cholinergic and noradrenergic activation support PF, similar to observations in the prefrontal cortex (PFC) but unlike noradrenergic suppression seen in the hippocampus. In contrast, dopamine and D1 receptor activation showed no clear effect on AC PF, contrasting with their suppressive role in the PFC. Furthermore, PF was significantly stronger in corticocollicular than corticocallosal neurons. These findings reveal diverse monoamine regulation of intrinsic PF, offering a potential cellular basis for sensory information maintenance under conditions of stress and reward.

## Introduction

Persistent neuronal firing, which is a repetitive spiking activity of neurons outlasting a triggering stimulus, has long been considered a fundamental cellular mechanism supporting short-term information storage, working memory, and context-dependent sensory processing (Fuster & Alexander, 1971; Goldman-Rakic, 1995; Major & Tank, 2004). Auditory cortex (AC) neurons exhibit persistent firing during tasks requiring auditory memory or attention across species. For instance, persistent firing in AC has been shown to reflect task-relevant information and support short-term retention in humans (Kumar et al., 2016). Delay-period persistent activity of AC neurons has been observed in nonhuman primates (Huang et al., 2016), rats (Sakurai, 1994) and mice (Yu et al., 2021), and correlates with auditory working memory (Huang et al., 2016; Yu et al., 2021). Persistent firing can arise from intrinsic membrane mechanisms, recurrent synaptic excitation, or a combination of both, often under the influence of neuromodulatory systems (Reboreda et al., 2018). Understanding how neuromodulators regulate persistent firing in defined cortical cell types remains essential for elucidating how internal brain states shape sensory and cognitive computations.

Acetylcholine (ACh) has been one of the most extensively studied neuromodulators in the context of persistent firing. Cholinergic signaling is strongly associated with arousal, attention, and learning (Hasselmo & Sarter, 2011), and numerous studies have shown that activation of muscarinic acetylcholine receptors can induce or enhance persistent firing in cortical and hippocampal pyramidal neurons (Egorov et al., 2002; Fraser & MacVicar, 1996; Klink & Alonso, 1997; Knauer et al., 2013; Navaroli et al., 2012; Tai et al., 2011). Mechanistically, cholinergic modulation engages calcium-dependent nonspecific cation currents and suppresses adaptation-promoting potassium conductances, thereby enabling sustained depolarization and spiking following transient input in individual cells (Haj-Dahmane & Andrade, 1998; Knauer & Yoshida, 2019; Tai et al., 2011). We refer to such persistent firing supported by individual neurons as “intrinsic persistent firing”. In the auditory cortex specifically, cholinergic terminals and receptor activation can induce persistent firing in layer V pyramidal neurons (Fu et al., 2019; Joshi et al., 2015), highlighting a potential cellular mechanism through which cholinergic inputs modulate auditory processing and memory-related functions.

Noradrenaline (NA, norepinephrine) is a neuromodulator released by the locus coeruleus and is strongly implicated in arousal, attention, and working memory (Aston-Jones & Cohen, 2005; Sara, 2009). In the prefrontal cortex, noradrenergic signaling enhances working memory performance by strengthening persistent firing and reducing noise in cortical networks (Arnsten & Goldman-Rakic, 1985; Wang et al., 2007). At the cellular level, NA can increase intrinsic excitability and promote sustained depolarizations by modulating potassium and calcium conductances (Constantinople & Bruno, 2011; McCormick et al., 1991). More recently, we and others have shown that NA supports persistent firing in prefrontal cortex cells while it suppresses persistent firing in the hippocampus (Valero-Aracama et al., 2021; Zhang et al., 2013). However, whether noradrenergic receptor activation can support persistent firing in auditory cortical pyramidal neurons, and how its effects compare to those of acetylcholine, remains largely unexplored.

Dopamine (DA) represents another key neuromodulator implicated in working memory in addition to reinforcement learning, motivation, and cognitive control (Schultz, 1998; Seamans & Yang, 2004). Particularly in the prefrontal cortex, dopaminergic modulation of persistent activity is essential for maintaining task-relevant information (Goldman-Rakic, 1995; Seamans & Yang, 2004). Both experimental and theoretical studies have demonstrated that DA can stabilize or destabilize persistent firing depending on receptor subtype and activation level, often following an inverted-U relationship (Durstewitz et al., 2000; Vijayraghavan et al., 2007). Both *in vivo* persistent firing and intrinsic persistent firing *in vitro* are suppressed by DA through D1/D5 receptor activation, suggesting a suppressive effect of DA in PFC and the entorhinal cortex (Batallán-Burrowes & Chapman, 2018; Lançon et al., 2021; Sidiropoulou et al., 2009; Vijayraghavan et al., 2007). In the auditory cortex, DA supports plasticity and learning where D1/D5 receptor activation enhances, and is required for, consolidation of auditory discrimination learning (Schicknick et al., 2008, 2012). It has also been shown that D1/D5 activation strengthens recurrent cortical feedback and promotes more sustained responses to relevant auditory stimuli, indicating DA can enhance persistent sensory representations (Happel et al., 2014). However, it remains unclear whether DA can directly support or modulate intrinsic persistent firing in auditory cortical neurons.

Cortical pyramidal neurons may also contribute to working memory in a projection-specific manner. In the prefrontal cortex (PFC), while pyramidal tract (PT) neurons show distinct contributions to working memory, intratelencephalic (IT) neurons provide temporal information (Bae et al., 2021). An *in vitro* study has indicated that intrinsic persistent firing is stronger in PT than IT neurons in the PFC (Dembrow et al., 2010). In the auditory cortex, corticocollicular (C-Col) neurons, which project to the inferior colliculus, and corticocallosal (C-Cal) neurons, which project to the contralateral cortex, represent two major and functionally distinct subpopulations (Slater et al., 2013). Previous work demonstrated that an activation of cholinergic terminals drives C-Col neurons to exhibit prolonged persistent firing, while C-Cal neurons did not (Joshi et al., 2016). However, whether this difference arises from intrinsic cellular properties and whether it depends on the type of neuromodulators has not been examined.

Together, existing studies support a central role for neuromodulators in enabling persistent firing associated with working memory, but leave unresolved how different neuromodulatory systems compare at the level of intrinsic cellular mechanisms of persistent firing in sensory cortex, and whether such modulations are distinct in comparison to executive and memory regions such as the PFC and the hippocampus. In particular, it remains unknown whether noradrenergic and dopaminergic signaling can support or modulate persistent firing in AC pyramidal neurons, and how such effects compare to cholinergic modulation. Furthermore, whether persistent firing is differentially invoked across distinct layer V projection neuron populations has not been addressed with respect to monoaminergic neuromodulation.

In the present study, we examined neuromodulator-supported persistent firing in layer V pyramidal neurons of the mouse auditory cortex by combining whole-cell patch-clamp recordings in acute brain slices with retrograde labeling and immunostaining to identify anatomically segregated projection targets and neuromodulatory influences. We compared the effects of cholinergic, noradrenergic, and dopaminergic receptor activation on persistent firing and assessed differences between corticocollicular and corticocallosal projection neurons. Our results demonstrate that cholinergic and noradrenergic, but not dopaminergic, receptor activation supports intrinsic persistent firing, and that this firing is significantly stronger in corticocollicular neurons. Combined with previous reports, these results indicate that cholinergic modulation exerts a similar supportive role for intrinsic persistent firing in the AC, PFC and the hippocampus. In contrast, the effect of NA on persistent firing in the AC differs from that in the hippocampus, while the effect of DA in the AC differs from that in the PFC. We further discuss the functional implications of these findings for maintaining sensory information under conditions of stress and reward. Together, these findings suggest that neuromodulation of intrinsic persistent firing may serve as a source of both state- and region-specific regulation of brain function.

## Methods

### In Vitro Patch Clamp Recording

All experimental protocols were approved by the Ethical Committee on Animal Health and Care of Saxony-Anhalt state, Germany (license number: 42502-2-1570 DZNE). 2-3 month C57BL6 male mice were deeply anesthetized with Isoflurane, assessed by the extinction of the pedal-withdrawal reflex, and decapitated. The brain was quickly removed from the skull and immersed in oxygenated ice-cold cutting solution containing (in mM) 110 Choline Cl, 7 MgCl2 (6H2O), 0.5 CaCl2, 2.5 KCl, 25 glucose, 1.2 NaH2PO4, 25 NaHCO3, 3 pyruvic acid, 11.5 ascorbic acid, and 100 D-Mannitol. Coronal brain slices (300 μm thick) were obtained with a vibrating-blade microtome (Leica VT1000S, Leica Biosystems, Wetzlar, Germany). Brain slices were then individually transferred to a holding chamber filled with normal ACSF (nACSF) containing (in mM) 126 NaCl, 1.2 NaH2PO4, 26 NaHCO3, 1.5 MgCl2, 1.6 CaCl2, 3 KCl, and 10 glucose. In this holding chamber, the slices were kept at 37 °C for 30 min and additionally for at least 30 min at room temperature before being used for recording. The pH of the cutting solution and nACSF were adjusted by a constant saturation with carbogen (95% O2/5% CO2).

Brain slices were transferred to the recording chamber and continuously superfused with nACSF (∼35 °C). The slice and cells were visualized with an upright microscope (Zeiss Axioskop 2FS plus, Carl Zeiss Microscopy, Jena, Germany) equipped with a 4× objective lens, a 40× water-immersion objective lens, and a monochrome camera (WAT-902H Ultimate). The auditory cortex was targeted using the (4x) objective lens, and a 40x objective was used to identify layer 5. Patch pipettes (3–8 MΩ) obtained from borosilicate glass capillaries (Science Products), were pulled on a P-1000 horizontal puller (Sutter instruments). The pipettes were filled with filtered intracellular solution containing (in mM): 120 K-gluconate, 10 HEPES, 0.2 EGTA, 20 KCl, 2 MgCl2, 7 PhCreat di(tris), 4 Na2ATP, and 0.3 Tris-GTP (pH adjusted to 7.3 with KOH). For labeling purposes, biocytin was added into the intracellular solution at 0.1% concentration. The whole-cell patch configuration was achieved by forming tight seals (>1 GΩ) on the soma and rupturing the membrane with light negative pressure. Liquid junction potential was not corrected. The access resistance was compensated several times during the recordings. Electrical signals were amplified with a Multiclamp 700B amplifier (Axon Instruments, Fremont, CA, USA), low-pass filtered at 10 kHz and sampled at 20 kHz, and recorded using pClamp (Axon Instruments, Fremont, CA, USA).

### Biocytin staining

To control for the anatomical location of recorded cells in layer V of the auditory cortex, biocytin staining was conducted. First, the brain slices were submerged in 4% PFA overnight before being transferred into 1X PBS. Within a month of being patched, the slices were transferred to a 24-well plate and washed 5x15 minutes in tris-buffered saline solution containing 0.5% Triton-X-100 (TBST-Tx, in mM: 153 NaCl, 50 Trizma Base, 0.5% Triton-X-100, pH adjusted to 8.0 using HCl).

After this, the slices were incubated overnight on a tilting mixer, covered or wrapped in foil to avoid light exposure, at room temperature with streptavidin-conjugated Alexa Fluor 647 (Thermo Fisher Scientific). The following day, the slices were rinsed three times for 15 min each in TBST-Tx, and mounted on standard microscope slides with Mowiol. Slides were stored overnight at 4° C, and imaged the next day using a Keyence (BZ-X710) fluorescent microscope.

### Measurements and data analysis

Measurement of persistent firing was done by depolarizing the cell to just below rheobase and applying a 2 s, 200 pA stimulus. Cells which continued firing for longer than three seconds after the end of the stimulus were considered to have persistent firing. Rheobase was determined prior to this protocol by slowly depolarizing a cell during a stimulus-free protocol and noting the point at which the cell fired an action potential. Persistent firing was classified as long-lasting if it continued longer than 30 s and short-lasting if it stopped before reaching 30 s. To quantify the strength of persistent firing, post-stimulus firing frequency and post-stimulus membrane potential were measured. Post-stimulus firing frequency was measured as the average firing rate of the cell during the 10 s period after the 2 s stimulus offset. Post-stimulus membrane potential was measured as the difference between the average membrane potential during the same 10 s period and the pre-stimulus baseline potential.

Intrinsic electrophysiological properties were measured during two protocols. In the first protocol, the resting membrane potential of the cell (V_rest_) was recorded without applying any current. After this, positive current was gradually applied until the cell was sufficiently depolarized to fire an action potential. Action potential amplitude, threshold, width at half-maximum, and maximum hyperpolarization speed were calculated from this action potential as described below using the scipy signal processing package (scipy.signal) in Python.

Action potential amplitude was measured as the difference between the voltage measured at the action potential’s peak and the average baseline measurement in the 50 ms directly before and after the spike. The width at half maximum was measured as the amount of time between the rising and falling phase of the action potential, measured at half of the action potential’s amplitude. The threshold of the action potential was calculated using the maximum of the third derivative of the voltage data. This corresponds to the point of the action potential where the voltage is changing fastest (Henze & Buzsáki, 2001). The maximum hyperpolarization speed was measured as the minimum of the first derivative of the voltage data, or where the voltage is decreasing most quickly.

In the second protocol (IV protocol), cells were held at a baseline membrane potential of -65 mV, and one-second-long square pulses were applied to the cell starting with -300 pA and increasing in 50 pA increments, ending with 400 pA. The input resistance was calculated by dividing the change in membrane potential resulting from the pulse by the amplitude of the pulse (-50 pA). Sag ratio was measured from the -300-pA pulse, and calculated by dividing the sag amplitude by the deflection of the membrane potential from baseline at peak sag current.

Burst index was measured from the first trace with at least 10 action potentials in the IV protocol. In most of the cells, we observed no burst or a burst with two spikes (doublet) at the initial part of the current injection. In these cases, the burst index was calculated as the second inter-spike-interval divided by the first inter-spike-interval. In some cells, more than two action potentials occurred in a burst riding on the same depolarizing wave at a rate of 60 Hz or faster. In those cases, the burst ratio was calculated as the first non-burst inter-spike-interval divided by the average inter-spike-interval during the burst.

Adaptation ratio was also measured from the same trace with at least 10 action potentials in the IV protocol. In cells that had either no burst or a burst with only two spikes (doublet), the adaptation ratio was calculated as the ratio of the 9th and 2nd inter-spike intervals. In cells with a burst with more than 2 spikes, the last non-burst and second non-burst inter-spike-interval are used instead. In rare cases, it was not possible to calculate the adjusted adaptation ratio on the first trace with 10 action potentials, in which case the next trace with 50 pA more current injection was used.

### Labeling of C-Col and C-Cal cells

In a subset of mice (n = 63), a retrograde tracer (100 nL green Lumafluor beads) was injected into either the contralateral primary auditory cortex (A1) or the ipsilateral inferior colliculus (IC) one week prior to patch-clamp recording. Mice were anesthetized with an intraperitoneal injection of a ketamine/xylazine mixture. After confirming the absence of a paw-pinch pedal reflex, the mice were fixed in a stereotaxic frame. Injection coordinates for the inferior colliculus (IC) were 1 mm posterior to lambda and 1 mm lateral, injection depth 0.75 mm (Joshi et al., 2015). Auditory cortex injection coordinates were 4 mm posterior to bregma and 4.5 mm lateral, injection depth 1 mm (Henschke et al., 2021). 100 nL of green Lumafluor retrograde tracing beads was injected through the craniotomy over the course of 10 minutes using a Hamilton syringe with a blunt metal needle (Hamilton 1701, I.D: 0.46 mm). The injection rate was controlled by a stereotaxic syringe pump (CHEMYX, NANOJET). After the injection was finished, the needle was left in place for an additional 5 minutes to allow the beads to disperse at the target location before being retracted.

The minimum time between injection surgery and patch clamp experiments was two days, in accordance with manufacturer instructions. However, the majority of mice were used 7-12 days after surgery. After the recording, Lumafluor beads, along with the biocytin-labelled cell, were visualized on a Keyence fluorescence microscope (BZ-X710).

### Classification of cell types using support vector machine

Based on the electrophysiological properties of labeled C-Col and C-Cal cells, we trained classifiers to assign non-labeled cells to these two groups. The classifiers were created in Python 3, using the scikit-learn library and tutorials. The following eight electrophysiological features from 60 C-Col and 62 C-Cal cells were used for the training: resistance, sag ratio, mAHP amplitude, V_rest_, maximum hyperpolarization speed, burst index, spike width at half maximum, sAHP amplitude, and adaptation ratio. Using the scikit-learn “pipeline” the features were first transformed to z-scores using the “StandardScaler” class, and then randomly divided into “train” (60%) and “test” (40%) groups. The training data was used to fit each classifier, and then the performance of the classifier was tested on the withheld 40% of labeled cells. We cross-validated the models using 10 random test-train sets to ensure that the outcomes were stable.

Based on performance and task suitability, linear SVM was chosen as the classification model and was implemented using the scikitlearn class “SVC” with a linear kernel. Because our classes were balanced, the “class-weight” parameter was not adjusted. The parameter “C” was reduced from the default setting (1) to 0.025, increasing the regularization to compensate for noise in the data and reduce the risk of overfitting. Other hyperparameters were not adjusted from the defaults provided by scikit-learn.

The classifier was evaluated using the F1 score, which measures how well the model identifies members of a class correctly (precision), without excluding members of that class (recall). The average F1 score over 10 cross-validation runs from the linear SVM classifier was 0.872 (SD = 0.077). Once trained, the model was “pickled” or serialized into a stable binary file, which could then be “unpickled” to run on a new dataset. In this way, the trained classifier was applied to unlabeled cells from other experiments. The model outputs a prediction of 1 or 2, corresponding to C-Cal and C-Col, respectively, along with additional information, such as the probability that the model assigned to each possible label.

All data analysis was performed using custom-written scripts in MATLAB (MathWorks) and Python.

## Results

### Cholinergic activation supports persistent firing in layer V AC cells

While the cholinergic agonist carbachol (Cch) has been widely used to induce persistent activity in a variety of brain areas, evidence from AC cells is limited (Fu et al., 2019). Therefore, before testing NA and DA, we first examined whether Cch supports persistent activity using a bath application of 5 µM Cch in deep-layer AC cells.

To evaluate the effects of Cch, we first tested persistent firing in the absence of Cch and then in its presence (5 µM). As in previous studies, persistent firing was tested from a membrane potential just below the spike threshold (Knauer et al., 2013). We refer to this as the baseline membrane potential; testing from this level enables us to compare the ability of cells to generate persistent firing independently of the varying resting membrane potentials and spike thresholds across cells. To trigger persistent firing, we used a brief current injection (100 pA, 2 s; Fig. 1A and C). In the control condition (without Cch), cells responded with spiking only during the current stimulation (n = 9).

**Figure 1.**
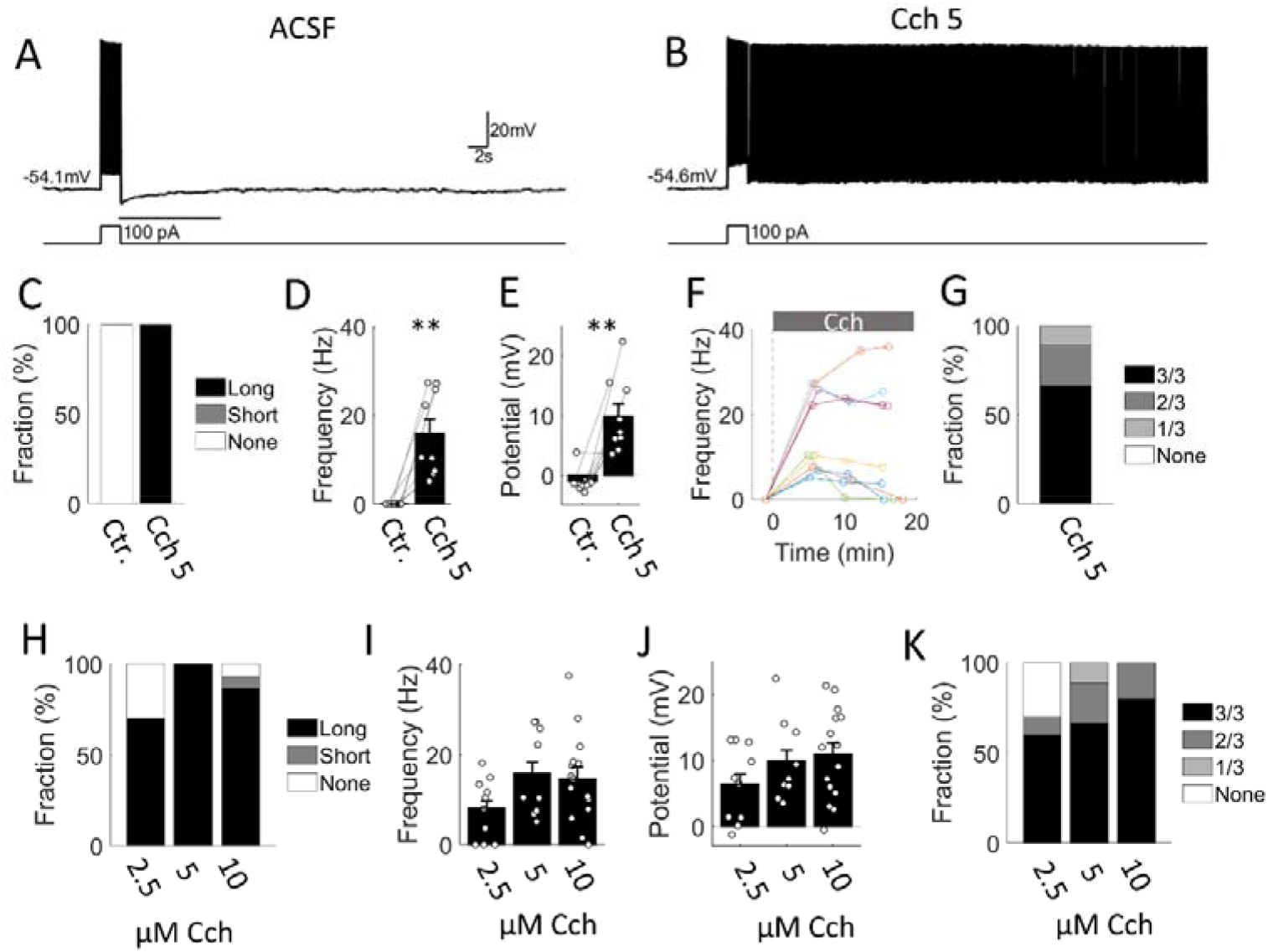
Cholinergic agonist carbachoi(Cch) supports persistent firing In layer V cells in the auditory cortex. (A and B) Example cell responding to a brief (2 s, 100pA) current stimulation in normal ACSF (control) and in 5 µM Cch. Upper trace shows the membrane voltage and the lower trace shows the current injection. The solid line between the voltage and current traces indicate the 10 s period from which the post-stimulus firing and membrane potential were measured. (C) Percentages of cells that responded with long, short and no persistent firing in control and Cch conditions (n = 9 and 9). (D) Post-stimulus firing frequency of cells in control and in 5 µM Cch (Wilcoxon signed-rank test; p = 0.004). (E) Post-stimulus membrane potential of cells in control and in 5 µM Cch (Wilcoxon signed-rank test; p = 0.008). (F) Post-stimulus firing rate of individual cells over time. Cch application started at time 0. (G) Response consistency (number of persistent firing response out of three attempts) in Cch. (H) Percentages of cells that responded with long, short and no persistent firing in different concentrations of Cch (n = 10,9,15). (I) Post-stimulus firing frequency of cells in different concentrations of Cch (Kruskal-Wallis test, H(2) = 3.45, p = 0.18). (J) Post-stimulus membrane potential of cells in different concentrations of Cch (Kruskal-Wallis test, H(2) = 2.24, p = 0.33). (K) Response consistency of cells in different concentrations of Cch.

Evaluation of persistent firing in the Cch condition was performed approximately 5 min after bath application of Cch in the same cells (n = 9). In contrast to the control condition, persistent firing was observed in all cells tested (Fig. 1B and C). Persistent firing was quantified by measuring the frequency and depolarization during the 10 s period immediately following the offset of the current injection, indicated by the line under the traces in Fig. 1A (see Methods for details of the quantification). The post-stimulus firing frequency, which ranged from ∼5 to 35 Hz, was significantly higher, and the membrane depolarization during the persistent firing phase was significantly larger in Cch compared to the control condition, indicating a supportive effect of Cch on persistent firing (Fig. 1D and E).

After this initial measurement, we repeated the protocol two more times in Cch to examine the consistency of persistent firing over time at approximately 10 and 15 min from the start of Cch application (Fig. 1F). The majority of cells (66.7 %) responded with persistent firing in all three attempts, while persistent firing was induced in two out of three attempts in the remaining cells (Fig. 1G).

We further tested persistent firing across a range of concentrations of Cch (2.5, 5, and 10 µM; Fig. 1H-K). The percentage of cells that responded with persistent firing was lower at 2.5 µM Cch compared to higher concentrations (Fig. 1H). The average frequency and depolarization of persistent firing increased from the lowest to the highest concentrations (Fig. 1I and J). However, there were no statistically significant differences among the concentrations. The response consistency also increased with increasing Cch concentrations (Fig. 1K).

### Projection of catecholaminergic fibers to the auditory cortex

Prior to testing the effects of NA and DA on persistent firing, we investigated whether NA and DA fibers project to the AC in mice. Immunohistochemical staining (Fig. 2A-C) revealed that fibers positive for tyrosine hydroxylase (TH), the rate-limiting enzyme for DA synthesis, are present in the AC (cyan stars in Fig. 2C top row, n = 3 mice). In addition, some TH+ fibers in the AC co-express DA β-hydroxylase (DBH), the enzyme responsible for converting DA to NA (pink triangles in Fig. 2C middle row, n = 2 mice). Together, these findings indicate that both DA- and NA-releasing fibers are present in the AC.

**Figure 2.**
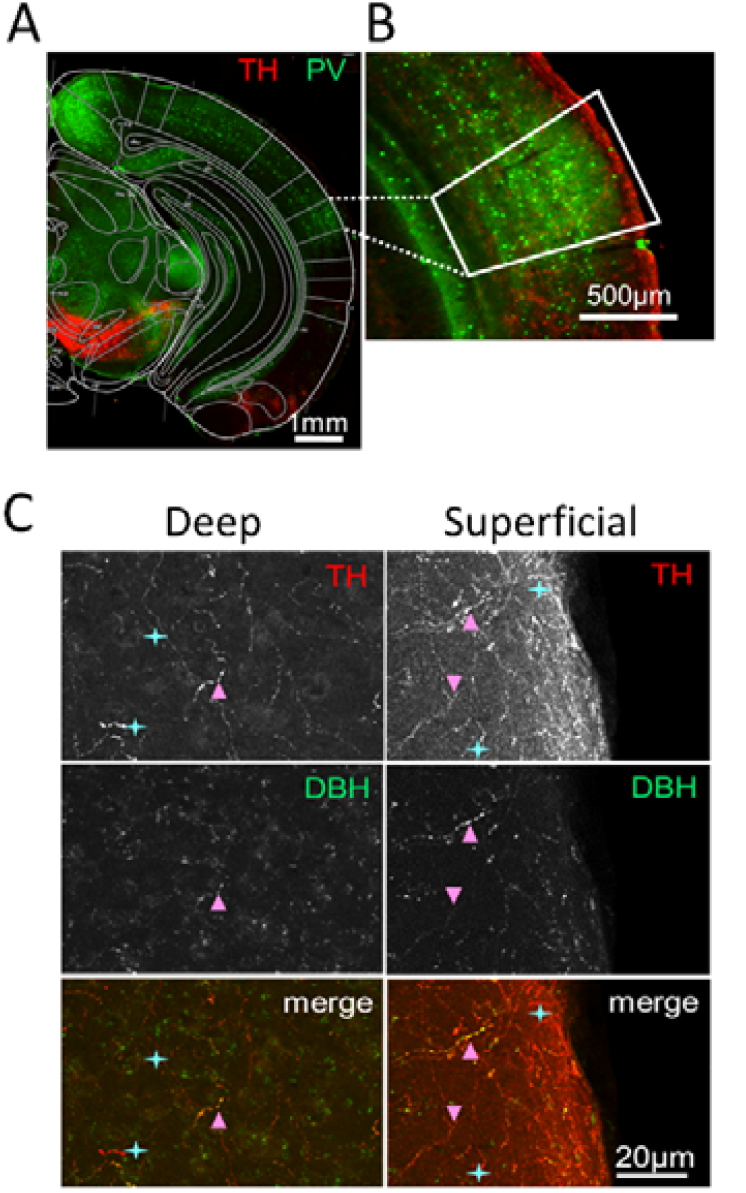
Presence of tyrosine hydroxylase (TH) and dopamine *β*-hydroxylase (DBH) positive fibers in auditory cortex. (A) Parvalbumin (PV) and TH staining. Intense PV staining indicates primary auditory cortex (Al). (B) Magnified Al area. (C) Double TH and DBH staining. Cyan stars: TH positive fibers. Pink triangles: DBH positive fibers.

### Noradrenergic activations support persistent firing

Next, we tested the effect of NA on the induction of persistent firing (n = 13). While persistent firing was not observed in the control condition, it was present in 54% of cells in 10 µM NA (Fig. 3A and B). The significantly higher firing frequency and membrane depolarization in NA relative to the control indicate a supportive effect of NA on persistent firing, similar to the action of Cch (Fig. 3C-E). Persistent firing was tested two additional times during the continued application of NA in each cell (Fig. 3F). At this concentration of NA, persistent firing was consistently observed (three out of three attempts) in only 38% of cells (5 out of 13), which was lower than the proportion observed in the Cch condition (Fig. 3G).

**Figure 3.**
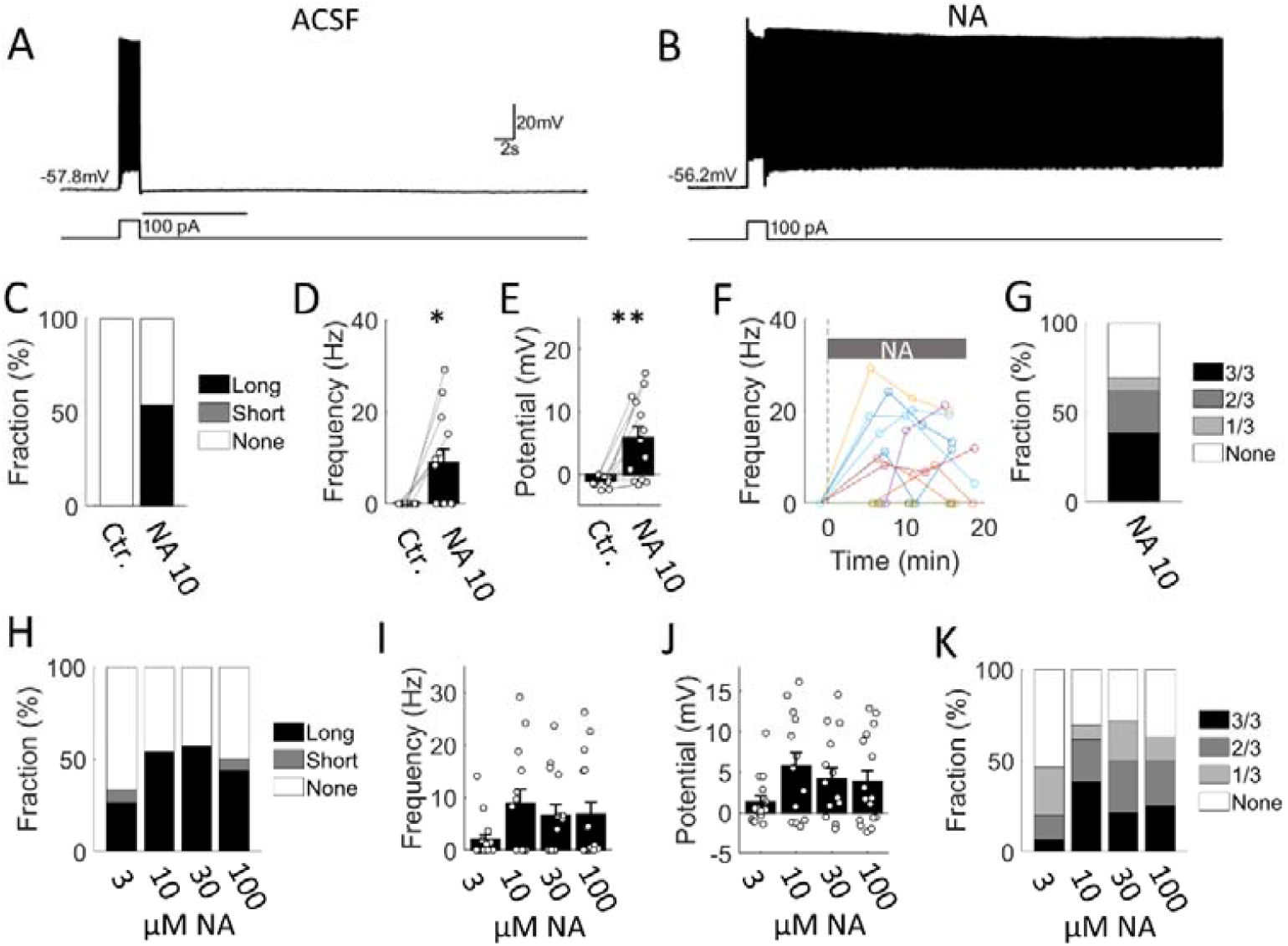
Noradrenaline (NA) supports persistent firing in layer V cells In the auditory cortex. (A and B) Example cell responding to a brief (2 s, 100pA) current stimulation in normal AC5F (control) and in 10 µM NA. Upper trace shows the membrane voltage and the lower trace shows the current injection. The solid line between the voltage and current traces indicate the 10 s period from which the post-stimulus firing and membrane potential were measured. (C) Percentages of cells that responded with long, short and no persistent firing in control and NA conditions (n = 13 and 13). (D) Post-stimulus firing frequency of cells in control and in 10 µM NA (Wilcoxon signed-rank test; p = 0.016). (E) Post-stimulus membrane potential of cells in control and in 10 µM NA (Wilcoxon signed-rank test; p = 0.002). (F) Post-stimulus firing rate of individual cells over time. NA application started at time 0. (G) Response consistency (number of persistent firing response out of three attempts) in NA. (H) Percentages of cells that responded with long, short and no persistent firing in different concentrations of NA (n = 15,13, 14,16). (I) Post-stimulus firing frequency of cells in different concentrations of NA (Kruskal-Wallis test, H(3) = 3.62, p = 0.31). (J) Post-stimulus membrane potential of cells in different concentrations of NA (Kruskal-Wallis test, H(3) = 2.71, p = 0.44). (K) Response consistency of cells in different concentrations of NA.

We further tested persistent firing across a range of concentrations of NA (3, 10, 30, and 100 µM). The percentage of cells that responded with persistent firing was lower at the lowest concentration (3.0 µM NA) compared to the higher concentrations (Fig. 3H). While average frequency and depolarization of persistent firing increased from the lowest to the higher concentrations (Fig. 3I), there were no statistically significant differences between them. The response consistency was lower in NA than in Cch even at the highest NA concentration (100 µM; Fig. 3K), suggesting that the relatively weaker effect of NA on persistent firing is not due to a lower potency or insufficient concentration of the drug.

### Cooperative action of NA and Cch on persistent activity

In *in vivo* conditions, both cholinergic and noradrenergic neuromodulation can be simultaneously active, potentially playing complementary roles in arousal, attention, and working memory. Therefore, we investigated whether NA and carbachol (Cch) can synergistically enhance persistent firing.

First, we recorded persistent firing in 10 µM NA and subsequently in a combination of 10 µM NA and 2.5 µM Cch in the same cells (n = 20, Fig. 4A and B). A higher percentage of cells exhibited persistent firing in the combined condition compared to the NA condition (Fig. 4C), and the persistent firing was, in general, stronger. The post-stimulus firing frequency was higher, and the depolarization was more pronounced in the combined condition (Fig. 4D and E).

**Figure 4.**
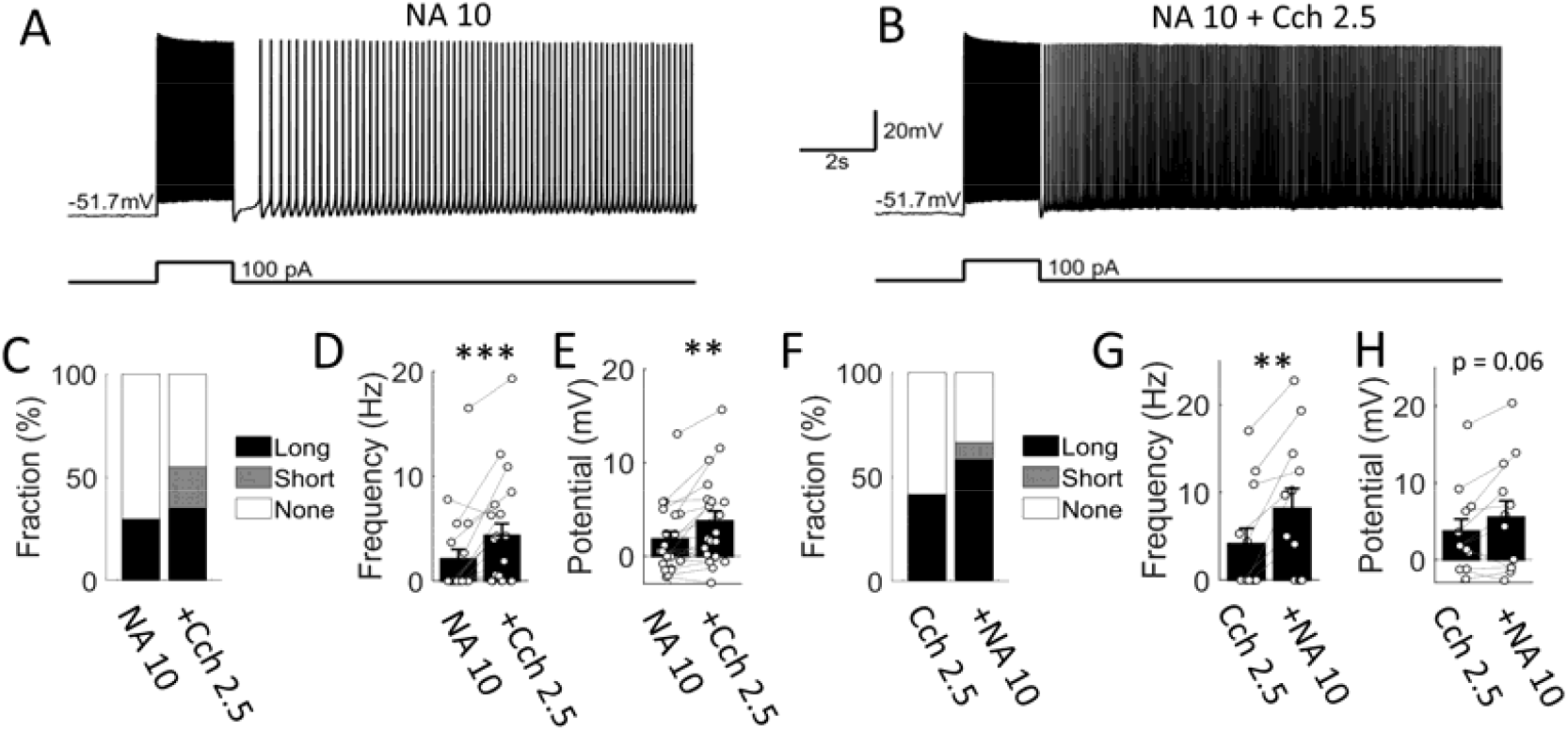
Cooperative action of NA and Cch on persistent activity. (A and B) Example cell responding to a brief (2 s, 100pA) current stimulation in 10 µm na and in the combination of 10 µM na and 2.5 µM Cch. (c) Percentages of cells that responded with long, short and no persistent firing in NA and in NA+Cch conditions (n = 20 for both). (D) Post stimulus firing frequency in NA and in NA+Cch conditions (Wilcoxon signed rank test, p < 0.001, n = 20). (E) Post-stimulus membrane potential in NA and in NA+Cch conditions (Wilcoxon signed-rank test, p = 0.0013, n = 20). (F) Percentages of cells that responded with long, short and no persistent tiring in 2.5 µM Cch and in the combination of 2.5 µM Cch and 10 µM NA conditions (n = 12 for both). (G) Post-stimulus firing frequency in Cch and Cch+NA conditions (Wilcoxon signed-rank test, p = 0.0078, n = 12). (H) Post-stimulus membrane potential in Cch and Cch+NA conditions (Wilcoxon signed-rank test, p = 0.064, n = 12).

We also tested the reverse application order. Persistent firing was initially tested in 2.5 µM Cch, followed by the addition of 10 µM NA to the same cells. The percentage of cells exhibiting persistent firing increased (Fig. 4F), and persistent firing was stronger in the combined condition (Fig. 4G). The depolarization showed a trend to be higher in the combined condition (Fig. 4H).

The cells used in this experiment came from a population of labeled C-Col and C-Cal cells. While comparison of persistent firing in these two groups of cells are done in Fig. 8 in Cch and NA conditions separately, we did not compare persistent firing in these two types of cells in the combined condition (Cch + NA) due to the limited number of cells tested. In summary, these results indicate that cholinergic and noradrenergic modulations are not occlusive and can act together to enhance intrinsic persistent firing.

### Dopamine does not strongly support or modulate persistent firing in auditory cortex

Dopaminergic modulation has been shown to improve working memory performance and enhance the signal-to-noise ratio of PFC activity by suppressing persistent firing in “off-target” cells (Vijayraghavan et al., 2007). The suppressive effect of DA on intrinsic persistent firing *in vitro* has also been shown in PFC cells (Lançon et al., 2021; Sidiropoulou et al., 2009). However, the effect of DA on intrinsic persistent firing in auditory cortex cells remains unknown.

While the effects of bath-applied DA on cells in the substantia nigra can be detected at concentrations as low as 100 nM and reach saturation at 10 µM (Philippart & Khaliq, 2018), other studies have used concentrations in the 30 µM to 50 µM range in the entorhinal (Caruana & Chapman, 2008) and prefrontal cortex (Otani et al., 2015). We therefore first tested the application of 30 µM DA in nine auditory cortex cells (Fig. 5A and B). Because DA oxidizes easily in solution, we used anti-oxidant sodium metabisulfite (50 µM), and stored the freshly prepared DA solution in low-light conditions. While none of the cells we tested responded with persistent firing in the control condition without DA, two of them responded with persistent firing ∼5 min after DA application (Fig. 5C). The frequency of persistent firing was not significantly different between the control and DA conditions (Fig. 5D), while the membrane potential following the stimulus increased slightly but significantly (Fig. 5E). This significant increase in membrane potential without persistent firing could reflect a decreased amplitude of the mAHP (Pedarzani & Storm, 1995). In seven of these nine cells, we tested persistent firing twice more over the following ∼15 min (Fig. 5F and G). The two cells that responded with persistent firing in the first test failed to do so in subsequent tests, suggesting an unreliable or transient effect of DA (Fig. 5F).

**Figure 5.**
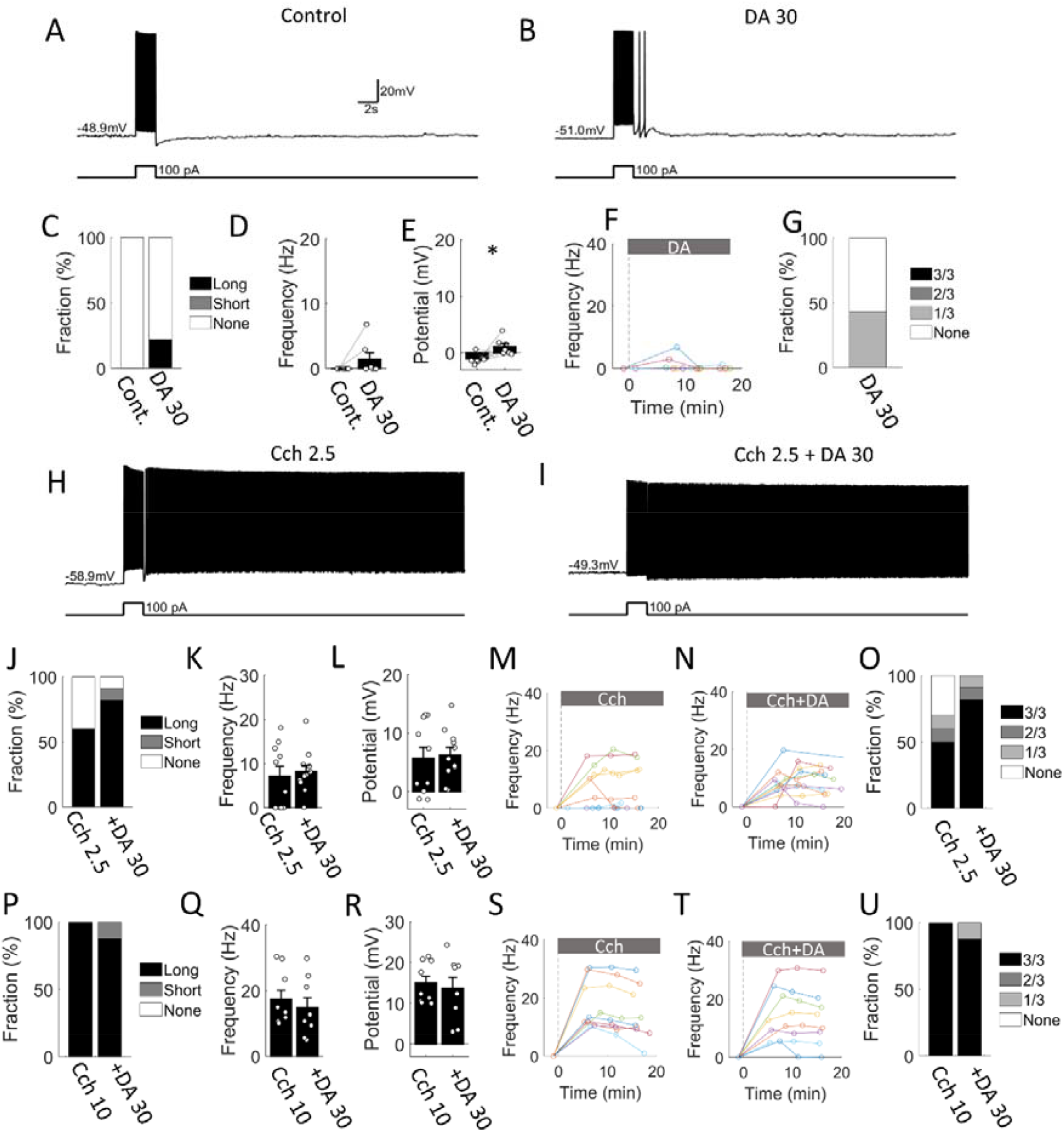
Dopamine (DA) does not strongly support or modulate persistent firing. (A and B) Example cell responding to a brief (2 s, 100pA) current stimulation in normal ACSF (control) and in 30 µM DA. (C) Percentages of cells that responded with long, short and no persistent firing in control and DA conditions (n = 9 for both). (D) Poststimulus firing frequency of cells in control and in NA conditions (Wilcoxon signed-rank test, p = 0.12, n = 9). (E) Post-stimulus membrane potential of cells in control and in DA (Wilcoxon signed-rank test, p = 0.012, n = 9). (F) Post-stimulus firing rate of individual cells over time. DA application started at time 0. (G) Response consistency (number of persistent firing response out of three attempts) in DA. (H and I) Example persistent firing in 2.5 µM Cch and in µM + 30 µM DA conditions. (J) Percentages of cells that responded with long, short and no persistent firing in Cch and Cch+DA conditions (n = 10 and 11). (K) Post-stimulus firing frequency of cells in Cch and Cch+DA conditions (Wilcoxon rank-sum test, p = 0.7). (L) Post-stimulus membrane potential of cells in Cch and Cch+DA conditions (Wilcoxon rank-sum test, p = 0.7). (M and N) Post-stimulus firing rate of individual cells over time. Drug application started at time 0. (O) Response consistency (number of persistent firing response out of three attempts) in Cch and Cch+DA conditions. (P-U) Same as J-0 but with 10 µM Cch (n = 9 and 8). Stats in Q: Wilcoxon rank-sum test, p = 0.42. Stats in R: Wilcoxon rank-sum test, p = 0.74 .

We next tested whether DA can modulate persistent firing in the presence of the cholinergic agonist Cch. Because we expected that DA might suppress Cch-induced persistent firing, and having observed that persistent firing gradually decreased over time in some cells during recording in Cch (Fig. 1F), we decided to compare persistent firing in Cch-alone and Cch+DA conditions in two different groups of cells at the same time point (∼5 min) after the start of drug application. Compared to the Cch (2.5 µM) condition, more cells responded with persistent firing in the Cch+DA (30 µM) condition (Fig. 5J; 2.5 µM Cch = 66.7% vs. 2.5 µM Cch + 30 µM DA = 100%). However, this effect was not statistically significant (Fisher’s exact test, p = 0.22). The frequency of persistent firing and post-stimulus depolarization did not differ significantly (Fig. 5K and L). Figs. 5M and N show measurements of persistent firing frequency from all cells at different time points (∼5, ∼10, and ∼15 min). The response consistency was higher in the Cch+DA condition (Fig. 5O).

The effect of DA on Cch-induced persistent firing was further assessed using a higher concentration of Cch (10 µM) in additional cells using the same experimental design (Fig. 5P-U). Neither the persistent firing frequency (Fig. 5Q) nor the post-stimulus depolarization level (Fig. 5R) differed between the two conditions. Overall, despite the innervation of the auditory cortex by tyrosine hydroxylase-positive fibers, the effects of DA on L5 pyramidal cells resulted in, at most, modest increases in post-stimulus depolarization, and we did not observe the suppressive effect expected based on results from the PFC.

### D1 receptor agonist does not suppress persistent firing

DA can activate both D1 and D2 receptor subtypes. The suppressive effect of DA on persistent firing *in vivo* in the PFC is mediated by the D1 receptor subtype (Vijayraghavan et al., 2007). This is in line with the fact that activation of D1 and downstream cAMP elevation suppresses intrinsic persistent firing *in vitro* in the PFC and the hippocampus (Lançon et al., 2021; Sidiropoulou et al., 2009; Valero-Aracama et al., 2021). D2 receptor activation, on the other hand, reduces cAMP, exerting the opposite effect of D1 activation (Beaulieu et al., 2015). To eliminate the possibility that our DA application above lacked a suppressive effect because the relatively high concentration of DA (30 µM) resulted in strong D2 activation that masked the action of D1 receptors (Trantham-Davidson et al., 2004), or that DA somehow failed to reach the cells, we tested the direct activation of D1 receptors using the D1 receptor agonist SKF81297 (1 µM and 10 µM).

The effect of SKF81297 was assessed in different groups of cells at a similar time point following drug application (∼15 min), consistent with the section above. We tested two different concentrations of SKF81297 (1 µM and 10 µM). When persistent firing was compared between cells that received Cch (10 µM) alone and Cch (10 µM) + SKF81297 (1 or 10 µM), there were no significant differences in either the firing frequency or depolarization (Fig. 6C-E). Similarly, comparisons performed at ∼30 min showed no differences among the three conditions, indicating that longer application of SKF does not exert different effects (Fig. 6F-H). These results suggest that D1 receptor activation does not suppress persistent firing in cells in the AC.

**Figure 6.**
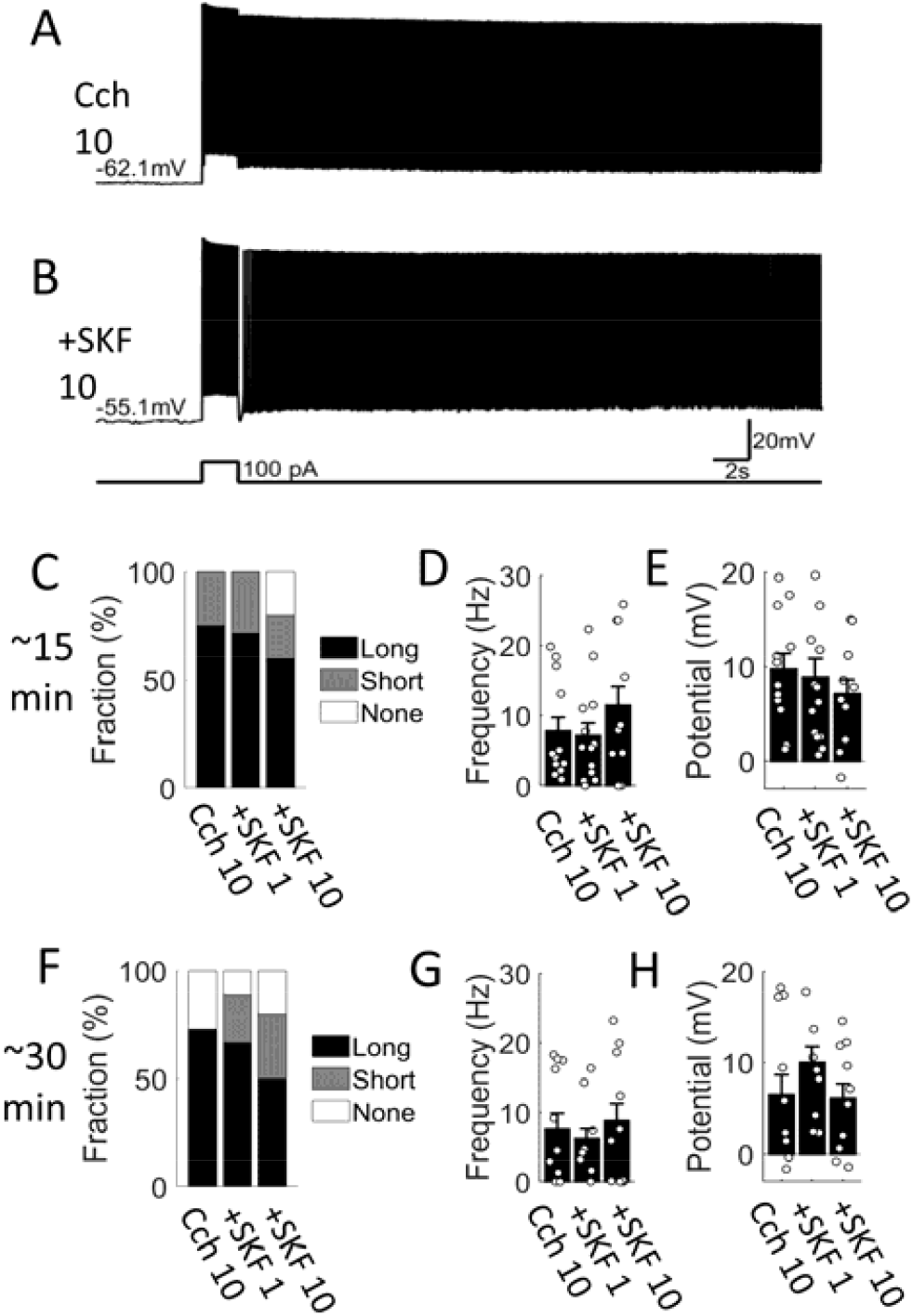
DI receptor agonist SKF81297 does not suppress persistent firing. (A and B) Example recordings of persistent firing in 10 pM Cch and in combination of 10 µM Cch and 10 µM SKF81297. (C) Percentages of cells that responded with long, short and no persistent firing in Cch and Cch+5KF81297 conditions (n = 12,14,10). (D) Post-stimulus firing frequency of cells in Cch and Cch+SKF81297 conditions (Kruskal-Wallis test, H(2) = 1.09, p = 0.58). (E) Post-stimulus membrane potential of cells in Cch and Cch+SKF81297 conditions (Kruskal-Wallis test, H(2) = 0.88, p = 0.65). (F-H) Same as C-E but using recordings from ∼30 min after drug application (n = 11,9, 10). Stats in G: Kruskal-Wallis test, H(2) = 0.28, p = 0.87. Stats in H: Kruskal-Wallis test, H(2) = 0.28, p = 0.87.

### Different strengths of persistent activity in C-Col and C-Cal cells

Cholinergically induced persistent firing *in vitro* is stronger in C-Col cells than in C-Cal cells (Joshi et al, 2016). However, it remains unknown whether the difference between C-Col and C-Cal cells arises from their intrinsic properties or from the different cholinergic projections they receive, because Joshi and colleagues used optogenetic stimulation of cholinergic fibers to induce PF. In addition, whether persistent firing supported by NA differs between these two types of cells remains unknown.

To identify C-Col vs. C-Cal neurons, we labeled cells using a retrograde tracer (100 nL green Lumafluor beads) injected into either the contralateral primary auditory cortex (A1) or the ipsilateral inferior colliculus (IC; Fig. 7A and B). As seen in Figs. 7C and D, C-Col cells were more likely to respond to a current injection with an initial burst of two or more action potentials. In addition, C-Col cells were characterized by a lower input resistance (Fig. 7E), a larger sag ratio (Fig. 7F), a more depolarized resting membrane potential (Fig. 7G), a larger spike frequency adaptation ratio (Fig. 7H), a higher burst index (Fig. 7I), and a narrower spike width (Fig. 7J). This replicates many of the differences previously reported (Joshi et al., 2015).

**Figure 7.**
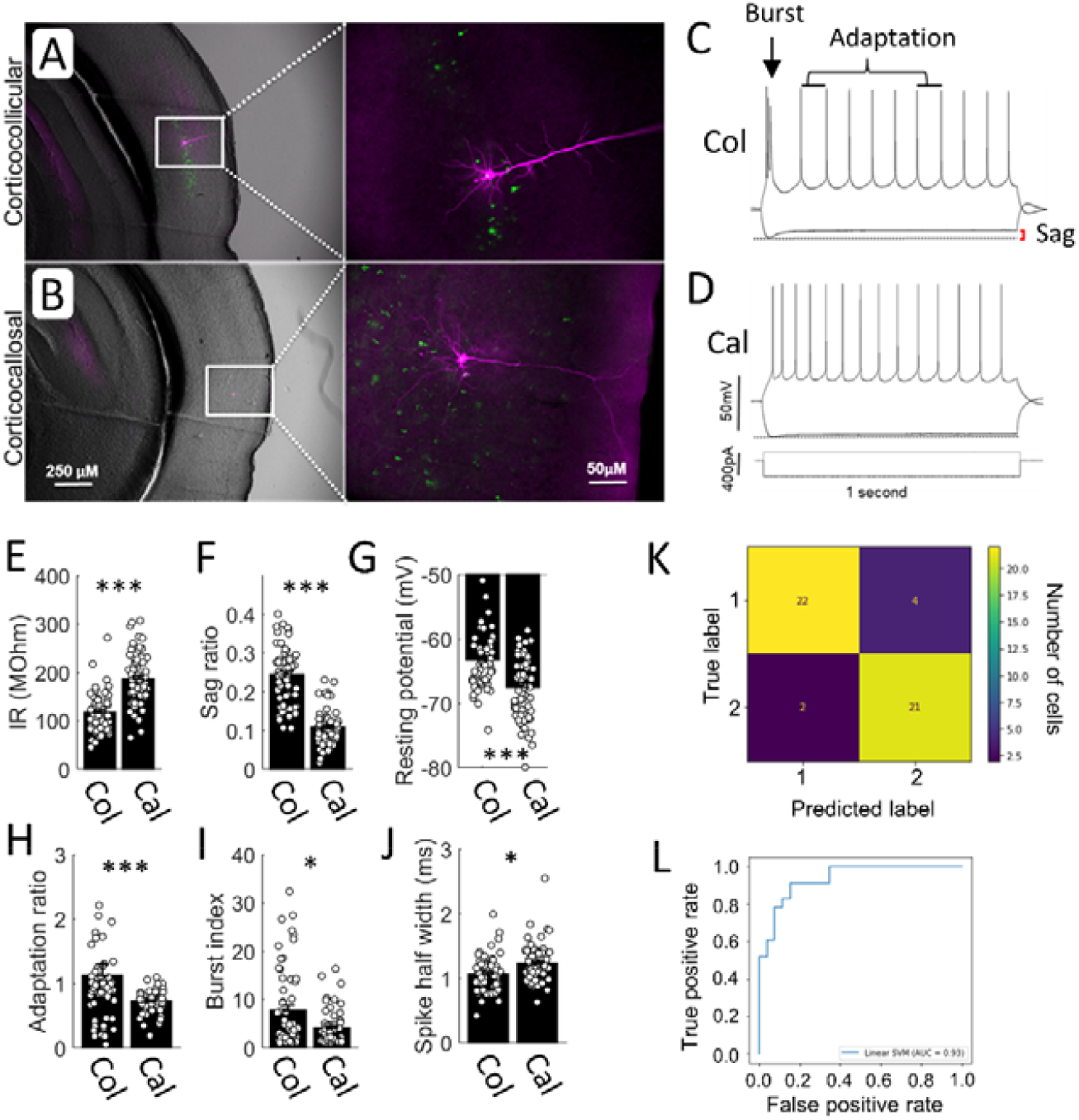
Classification of corticocollicular (C-Col) and corticocallosal (C-Cal) cells. (A and B) Examples of labeled C-Col and C-Cal cells using retrograde tracer (green Lumafluor beads). Recorded cell is stained using biocytin and is shown as magenta. (C and D) Membrane potential response to negative and positive square pulse from example C-Col and C-Cal cells. (E) Input resistance (n = 60 and 62; Two-tailed T-test, p < 0.001). (F) Sag ratio (n = 60 and 62; Wilcoxon rank-sum test, p < 0.001). (G) Resting membrane potential (n = 60 and 62; Wilcoxon rank sum test, p < 0.001). (H) Spike frequency adaptation ratio (n = 60 and 62; Wilcoxon rank-sum test, p < 0.001). (I) Burst index (n = 60 and 62; Wilcoxon ranksum test, p = 0.023). (J) Spike half width (n = 60 and 62; Wilcoxon rank sum test, p = 0.0028). (K) Example confusion matrix for Linear SVM. Label 1 and 2 correspond to C-Cal and C-Col, respectively. (L) Receiver operating characteristic (ROC) showing the proportion of true positives (hits) and false negatives (misses) identified at a given false positive rate. ROC area under curve (AUC) = 0.93.

While these features are significantly different between the two groups, there is no single feature that can be used to distinguish the two cell types. Therefore, we built a Support Vector Machine (SVM) classifier to classify unlabeled cells based on eight different electrophysiological properties (Fig. 7K and L; See Methods). This classifier could accurately classify labeled cells with an F1-score of 0.872 (SD = 0.077; Fig. 7K and L) and was used to compare persistent firing in C-Col and C-Cal cells below.

Persistent firing in C-Col and C-Cal cells was first compared in 10 µM Cch using only labeled cells (10 C-Col and 11 C-Cal cells; Fig. 8A and B). In each cell, persistent firing was measured at approximately 5 min following the application of Cch (∼15 min from break-in). While almost all (90%; 9/10) of the C-Col cells exhibited persistent firing, a lower percentage (63.6%; 7/11) of C-Cal cells did so (Fig. 8C). The firing rate and depolarization of persistent firing were both significantly higher in C-Col cells than in C-Cal cells (Fig. 8D and E). In Figs. 8F to H, we compared persistent firing by combining cells that were labeled and classified by SVM (19 C-Col and 17 C-Cal cells). As expected, the frequency and depolarization of persistent firing were higher in the C-Col cells (Fig. 8F-H).

**Figure 8.**
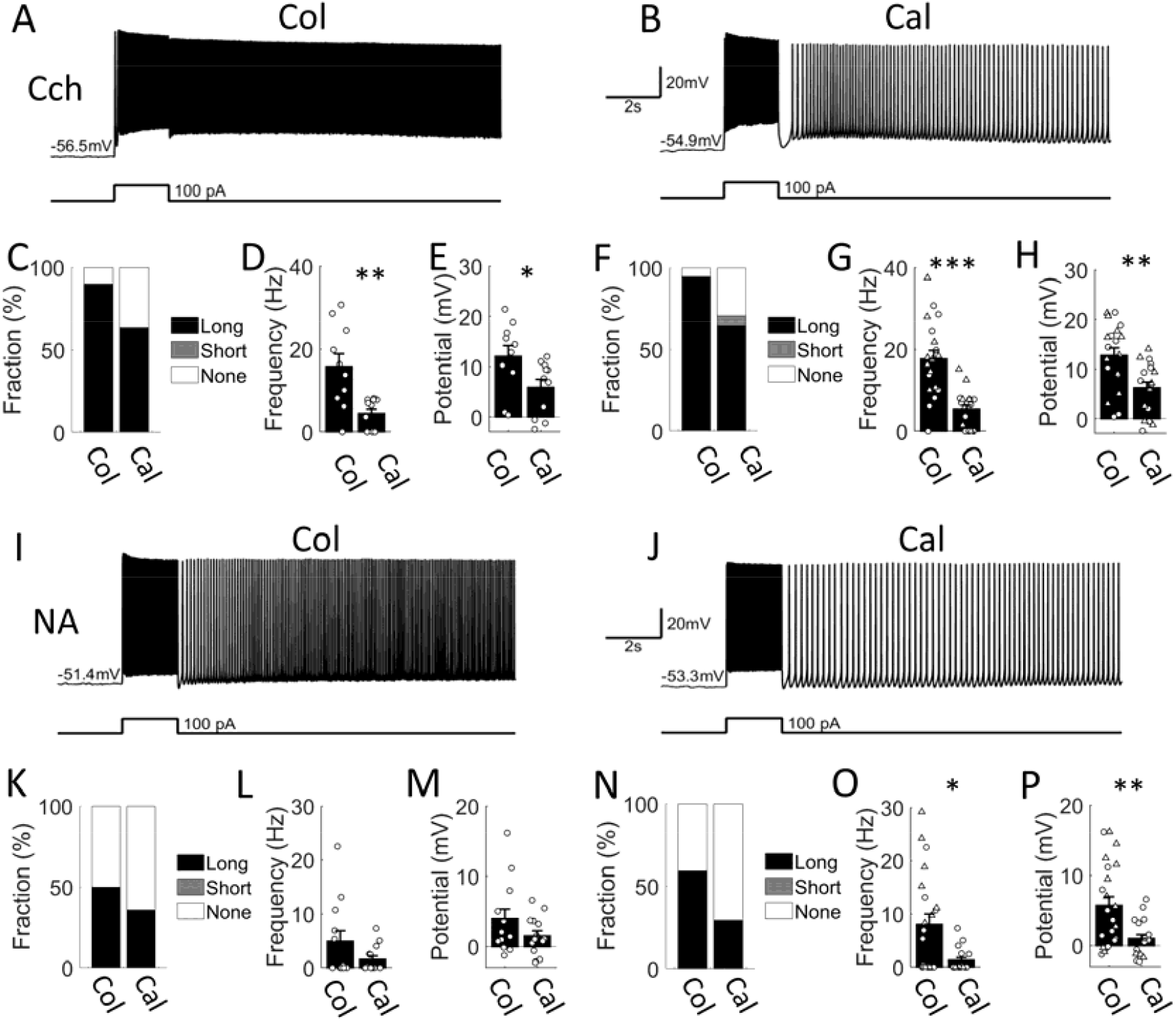
Persistent firing is stronger in C-Col than in C-Cal cells both in Cch and NA. (A and B) Examples of persistent firing in a labeled C-Col and C-Cal cells In 10 µM Cch. (C) Percentages of labeled C-Col and C-Cal cells that responded with long, short and no persistent firing in 10 µM Cch (n = 10 and 11). (D) Poststimulus firing frequency of labeled C-Col and C-Cal cells in 10 µM Cch (Wilcoxon rank-sum test; p = 0.005). (E) Post-stimulus membrane potential of labeled C-Col and C-Cal cells in 10 µM Cch (Wilcoxon rank-sum test; p = 0.031). (F-H) Same as C-F but fnr combined population of labeled and classified cells (n = 19 and 17; Wilcoxon rank-sum test; p < 0.001 and p = 0.003 in G and H, respectively). (I and J) Examples of persistent firing in a labeled C-Col and C-Cal cells in 10 µM NA. (K) Percentages of labeled C-Col and C-Cal cells that responded with long, short and no persistent tiring in 10 µM NA (n = 12 and 14). (L) Post-stimulus firing frequency of labeled C-Col and C-Cal cells in 10 µM NA (Wilcoxon rank-sum test; p = 0.36). (M) Poststimulus membrane potential of labeled C-Col and C-Cal cells in 10 µM NA (Wilcoxon rank-sum test; p = 0.34). (N-P) Same as K-M but for combined population of labeled and classified cells (n = 22 and 17; Wilcoxon rank-sum test; p = 0.021 and 0.008 in O and P, respectively).

To test if this higher frequency of persistent firing in C-Col cells resulted only from a higher percentage of cells responding with persistent firing or not, we also compared persistent firing only in cells that responded with persistent firing (18 C-Col and 12 C-Cal cells). We found that persistent firing was still significantly stronger in C-Col cells (18.60 ± 2.02 Hz) than in C-Cal cells (7.63 ± 1.03 Hz), indicating that when C-Col cells respond with persistent firing, the activity is stronger than in C-Cal cells (Wilcoxon rank-sum test, p < 0.001; data not shown). These results suggest that C-Col cells are more prone to exhibiting persistent firing and the intensity of the induced persistent firing is stronger. This extends previous reports (Joshi et al., 2016) by indicating that stronger persistent firing in C-Col cells originates from mechanisms intrinsic to the individual cells.

Similar comparisons were made in 10 µM NA as well (Fig. 8I to P). While the strength of persistent firing in labeled cells did not reach statistical significance (12 C-Col and 14 C-Cal cells; Fig. 8K-M), an analysis combining labeled and classified cells indicated that persistent firing in NA is also stronger in C-Col cells than in C-Cal cells (22 C-Col and 17 C-Cal cells; Fig. 8N-P). Similar to the case with Cch, persistent firing was stronger in C-Col cells (13.49 ± 2.28 Hz) than in C-Cal cells (3.57 ± 1.00 Hz) when the comparison was restricted to cells exhibiting persistent firing (Wilcoxon rank-sum test, p = 0.0032). These results suggest that persistent firing is stronger in C-Col cells than in C-Cal cells, not only under cholinergic but also under noradrenergic neuromodulation.

## Discussion

In this study, we investigated the neuromodulatory control of intrinsic persistent firing in layer V pyramidal neurons of the mouse auditory cortex. Our results revealed that while cholinergic and noradrenergic activation robustly support intrinsic persistent firing, dopaminergic signaling does not. Furthermore, we demonstrated that this capacity for sustained activity is non-uniformly distributed across cortical output pathways, with corticocollicular (C-Col) neurons exhibiting significantly stronger persistent firing than corticocallosal (C-Cal) neurons under both cholinergic and noradrenergic modulations.

### Differential roles of NA on intrinsic persistent firing among different areas

Our results establish NA as a potent driver of intrinsic persistent firing in layer V auditory cortex neurons, adding to a growing body of evidence that NA acts as an important modulator of sustained activity. This finding is consistent with observations in the prefrontal cortex, where NA has been shown to drive persistent firing through the synergistic activation of ⍰1 and ⍰2 adrenoceptors (Zhang et al., 2013). The robust induction of persistent activity we observed in the auditory cortex suggests that NA serves to stabilize and prolong internal representations during states of arousal in both sensory and executive cortices. However, this effect stands in stark contrast to recent findings in the hippocampus, where NA was found to suppress intrinsic persistent firing in CA1 pyramidal neurons through β1 and β2 adrenergic receptors (Valero-Aracama et al., 2021). While the functional role of this divergence between the cortex and the hippocampus requires further study, it is interesting to speculate that high levels of in vivo NA associated with acute stress or high arousal prioritize the processing of immediate external threats over the detailed encoding of contextual or episodic information in the hippocampus.

### Potential difference of actions of NA and Cch on persistent firing

While NA induced persistent firing in the majority of cells tested, the percentage of cells that responded with persistent firing and the average frequency of persistent firing were lower than in Cch (Fig. 2). This could be due to the activation of β-adrenergic receptors that trigger Gs-coupled signaling, elevate cAMP/PKA, and suppress the TRPC-mediated Ca^2+^-activated nonselective cation (CAN) current required for a stable depolarizing plateau (Reboreda et al., 2018). In contrast, cholinergic modulation does not activate the Gs-pathway, whereas it activates the Gq-pathway through M1 and M3 receptors, and the Gi-pathway through M2 and M4. These pathways are well known to engage the TRPC/CAN current and suppress the M-current, thereby promoting robust persistent firing (Reboreda et al., 2018).

### Action of DA on intrinsic persistent firing

The DA application recruited persistent firing in a subset of cells we recorded. However, this effect was not as strong or reliable as those of NA and Cch. On the other hand, we observed a small but significant increase in the post-stimulus membrane potential in DA. This is in line with the action of DA in the PFC to reduce the spike after-hyperpolarization potential (AHP; (Thurley et al., 2008), and it also indicates that DA was acting on the cells recorded. While the currents underlying the AHP, such as the M-current or SK-current, are different from the CAN current that directly drives persistent firing, AHP currents do modulate persistent firing. We have shown that M-current suppression, which happens under cholinergic activation, uncovers persistent firing in the MEC and strengthens it in hippocampal neurons (Knauer & Yoshida, 2019; Yoshida & Alonso, 2007). Furthermore, a recent study has shown that suppression of the M-current contributes to persistent firing in auditory cortex layer V neurons (Ye et al., 2022).

While the reduction of the AHP increases intrinsic excitability, studies done in the PFC and entorhinal cortex (EC) have shown a clear suppressive effect of DA on persistent firing and plateau depolarization through the D1 receptor subtype (Batallán-Burrowes & Chapman, 2018; Lançon et al., 2021; Sidiropoulou et al., 2009). Therefore, we further tested the application of DA or a D1-receptor agonist (SKF81297) on top of Cch to specifically test whether DA and D1 activation can suppress persistent firing. To our surprise, we did not observe a suppressive effect of DA or SKF81297 on Cch-induced persistent firing, suggesting that, similarly to the action of NA, DA signaling is also region-specific. A speculative functional implication of this could be that while reward-related DA elevation could reset PFC-mediated goal-directed behavior for task-switching, sensory cortices are spared from this suppression, allowing them to continue environmental monitoring by maintaining sensory representations.

### Persistent firing in C-Col vs C-Cal

By using bath application of Cch and NA, we extend the findings of Joshi et al. (2016), who reported that C-Col cells respond with a stronger persistent firing when cholinergic fiber activation is used. While cholinergic fiber activation is a more physiological way to trigger persistent firing, the observed difference could stem from both potential differences in cholinergic fiber innervation and the intrinsic cellular mechanisms of these cell types. Using a controlled strength of cholinergic activation, our data suggest that the differences arise from intrinsic cellular mechanisms.

This aligns with and extends the emerging “projection-specific” model of cortical function. In the prefrontal cortex (PFC), a similar dichotomy exists: pyramidal tract (PT) neurons, which project to subcortical structures like the pons or medulla, exhibit robust, stable persistent firing, whereas intratelencephalic (IT) neurons, which project to the contralateral cortex or striatum, show transient or weaker sustained activity (Bae et al., 2021; Dembrow et al., 2010). Since C-Col neurons are a subset of the PT population and C-Cal neurons belong to the IT population, our data suggest that the cellular machinery for persistent firing is a conserved feature of PT-type neurons across different functional cortical areas.

### Intrinsic vs. synaptic origin of persistent firing

Finally, while the literature often emphasizes the role of recurrent synaptic loops in maintaining activity, recent studies indicate the potential involvement of an intrinsic origin for persistent firing. Manipulation of the molecular mechanisms of intrinsic persistent firing, such as the TRPC channels and p75NTR, affects working memory (Bröker-Lai et al., 2017; Gibon et al., 2015; Lepannetier et al., 2018). Intrinsic persistent firing *in vitro* is stronger in animals that successfully acquired trace eyeblink conditioning (Lin et al., 2020). In addition, we recently reported that TRPC4 knockdown (KD), which disrupts intrinsic persistent firing in the hippocampal cells, indeed reduces persistent firing in vivo, providing the first direct evidence that intrinsic persistent firing supports in vivo persistent firing during a working memory task (Saber Marouf et al., 2025). Future work should focus on whether differential modulation of in vivo persistent firing by neuromodulators depends on the differential modulation of intrinsic persistent firing, and whether differences of persistent firing in vivo in C-Col (PT) and C-Cal (IT) cells reflect the intrinsic cellular properties studied here.

## Acknowledgments

We would like to thank C. Knape and J. Henschke for their help in establishing injection surgery procedures, to Y. Huang and M. Brosch for useful discussions, and to M. Lippert, C. Helbing and M. Sauvage for technical support. This work was supported by Center for Behavioral Brain Sciences (CBBS) NeuroNetwork program, and the Deutsche Forschungsgemeinschaft projects YO177/4-1, YO177/4-3, and YO177/7-1 (to M.Y.).

## Notes

### Competing Interest Statement

The authors have declared no competing interest.

